# Accessible and Robust Machine Learning Approaches to Improve the Opsin Genotype-Phenotype Map

**DOI:** 10.1101/2025.08.22.671864

**Authors:** Seth A. Frazer, Todd H. Oakley

## Abstract

Predicting phenotypes from genetic variation is a central challenge in biology. Linking genotypes and phenotypes using machine learning (ML) offers great promise, but its use is limited by poor accessibility, overestimated performance, and a “data-cliff”—a gap between abundant sequences and scarce functional measurements. To develop more robust methods for genotype–phenotype prediction, an outstanding model system is opsin genes, visual pigments with extensive phenotypic information that strongly influence animal spectral sensitivity. Here we advance ML characterization of the opsin genotype–phenotype map through four main contributions. First, we introduce the *Opsin Phenotype Tool for Inference of Color Sensitivity* (OPTICS), a user-friendly platform for predicting maximum wavelength sensitivity (λ_max_) from amino-acid sequences. Second, we show that encoding sequences with amino-acid physicochemical properties improves predictive performance and reveals mechanistic relationships. Third, we develop Phylogenetically Weighted Cross-Validation (PW-CV), a method that accounts for non-independence among related sequences, providing more realistic assessments of model generalizability. Finally, we present the Mine-N-Match (MNM) pipeline, which systematically links published opsin sequences to compiled in-vivo λ_max_ data, expanding genotype–phenotype coverage and improving prediction, especially for invertebrate opsins with undersampled heterologous data. By integrating accessible software, biologically informed encoding, phylogeny-aware evaluation, and data harmonization, our framework improves confidence, accuracy, and interpretability of genotype–phenotype prediction. An accurate genotype-phenotype map allows simulating molecular evolution of function, reconstructing the history of visual phenotypes, designing functional proteins, and generating new hypotheses that can be tested with heterologous phenotyping.

## Introduction

Accurately relating phenotypes and genetic variation is a central challenge in evolutionary biology (Wagner 2011; Kemble et al. 2019; Nichol et al. 2019). A strong understanding of such genotype-phenotype maps will allow reconstructing historical shifts in gene function (Yokoyama et al. 2014; Storz 2016) and simulating or forecasting evolutionary pathways (de Visser and Krug 2014; Lässig et al. 2017). Accurate genotype-phenotype maps could transform fields from molecular evolution to medicine, agriculture, and protein engineering. Yet despite recent progress in heterologous expression assays (Liénard et al. 2022; Smedley et al. 2022) and high-throughput mutational scanning (Fowler and Fields 2014; Inoue et al. 2021), empirical approaches remain costly and time-consuming, limiting their scalability for broad comparative studies.

Artificial intelligence (AI) including Machine learning (ML) provides a powerful complement to empirical work by learning predictive rules from genotype–phenotype datasets. These computational approaches accelerate discovery, reduce dependence on labor-intensive empirical assays, and generate testable hypotheses for experimental validation (Vamathevan et al. 2019; Hie et al. 2020; Frazer et al. 2021). For example, protein language models trained on millions of sequences capture structural and functional constraints (Rives et al. 2021; Lin et al. 2023) and large language models (LLMs) can simulate protein evolution and forecast mutational effects across deep timescales (Hayes et al. 2025). Despite this promise, barriers remain. First, most ML/AI tools require computational expertise, limiting accessibility to experimental biologists. Second, commonly used evaluation schemes often overestimate predictive performance by ignoring phylogenetic non-independence among sequences (Roberts et al. 2017). Third, a persistent “data-cliff”—a steep imbalance between abundant sequences and scarce functional measurements—limits the training of robust, generalizable models (Bhardwaj et al. 2025). Overcoming these barriers is essential if genotype–phenotype prediction is to become a reliable, widely used tool in molecular evolution.

To address these barriers, opsins provide an outstanding system for advancing genotype–phenotype prediction (Karasuyama et al. 2018; Adam et al. 2022; Frazer et al. 2024; Sela et al. 2024). These G-protein Coupled Receptors form the molecular basis of animal vision and determine color sensitivity through their maximum wavelength of light absorption (λ_max_) (Govardovskii et al. 2000). Opsins are especially well-suited for methodological development because they combine massive sequence diversity with extensive phenotypic data from heterologous expression, in vivo measurements, and comparative studies (Terakita 2005; Yokoyama 2008; Hagen et al. 2023). Moreover, opsins have repeatedly diversified across metazoans, offering natural experiments in functional convergence and divergence (Porter et al. 2012; Ramirez et al. 2016; Vöcking et al. 2017). This combination of broad sequence sampling, rich functional characterization, and evolutionary replicates makes opsins an excellent model for testing and refining ML approaches to genotype–phenotype mapping.

Here we advance machine learning modeling of the opsin genotype–phenotype map through four main contributions. First, we introduce the *Opsin Phenotype Tool for Inference of Color Sensitivity* (OPTICS), a user-friendly platform that predicts maximum wavelength sensitivity (λ_max_) from amino-acid sequences. Second, we show that encoding sequences with physicochemical properties of amino-acids improves predictive accuracy and enhances mechanistic interpretability. Third, we develop Phylogenetically Weighted Cross-Validation (PW-CV), a method that accounts for non-independence among related sequences, yielding more realistic assessments of model generalizability. Finally, we present the *Mine-N-Match* (MNM) pipeline, which systematically links published opsin sequences to compiled in vivo λ_max_data, expanding genotype–phenotype coverage and improving prediction, especially for invertebrate opsins with sparse heterologous expression data. Together, these advances establish a methodological foundation for predictive molecular evolution using opsins as a model system—opening the door to simulating molecular evolution, reconstructing ancestral visual phenotypes, and generating testable hypotheses for experimental validation.

## Methods

### OPTICS: An accessible ML tool for predicting opsin spectral sensitivity

To improve the accessibility of our machine learning (ML) models, we developed the *Opsin Phenotype Tool for Inference of Color Sensitivity* (OPTICS), a user-friendly tool for predicting opsin λ_max_ values from unaligned amino-acid sequences. OPTICS is available as a command-line tool, graphical-user-interface (GUI), and a web application under the Galaxy system; accessible through GitHub (github.com/VisualPhysiologyDB/optics). OPTICS provides users eleven best-performing ML models, trained on eleven different datasets, which can use either one-hot encoding (scoring each amino acid at each position as either present or absent) or an ‘amino-acid property’ (aa-property) encoding scheme described herein. For prediction, OPTICS aligns the user’s query sequence to the training alignment of the selected model using MAFFT (Katoh and Standley 2013).

In addition, OPTICS incorporates optional bootstrapping and BLASTp (Camacho et al. 2009) analyses. Bootstrapping (described in more detail below) quantifies uncertainty, including box plots, colored to reflect the median predicted λ_max_ using the corresponding hex-code color value. The BLASTp analysis compares query sequences against the training dataset of the selected model. Site numbering in BLASTp reports can be standardized to a chosen reference sequence (e.g., bovine or squid rhodopsin) or a custom one (see Supplemental Methods S1 for implementation details and availability). OPTICS is implemented in Python, with core functionality built using scikit-learn (Pedregosa et al. 2011) for model execution and BioPython (Cock et al. 2009) for sequence handling, along with dependencies such as MAFFT, BLAST+, and standard Python data-processing libraries.

### Assessing prediction stability with bootstrapping

To evaluate the stability of λ_max_ predictions and estimate confidence intervals, we implemented a bootstrapping pipeline. For each VPOD opsin data subset (Frazer et al. 2024), we generated 100 pseudoreplicated datasets by randomly resampling with replacement entire sequences and their associated λ_max_ value. For each pseudoreplicate we trained a separate ML model for predicting λ_max_ for sequences input into OPTICS. We report mean and median predictions across replicates, using the standard deviation to construct confidence intervals (see Supplemental Methods S2). To illustrate the procedure, we applied bootstrapping to three example sequences with disparate λ_max_: a short-wave sensitive (SW) opsin (OX393528.1-9544021-9545976) from the Tobacco Hornworm (*Manduca sexta*), a rhodopsin (Rho; MN519143.1) from the Bluespotted Stingray (*Neotrygon kuhlii*), and a long-wave sensitive (LWS) opsin (MK983124.1) from the Corkwing Wrasse (*Symphodus melops*).

### Encoding amino acid physicochemical properties for ML model training

We compared traditional one-hot encoding (each amino acid scored as present or absent at each position) to a novel encoding based on twelve physicochemical properties (aa-properties; Supplemental Methods S3, Supplementary Table). In the new approach, we represented each amino acid as continuous values of amino acid properties. Unlike one-hot encoding which treats all unseen residues at a site as equivalent, aa-property encoding allows all models to use information for residues not present in the training set (Supplementary Fig. 1B). We used *deepBreaks* (Baghbanzadeh et al. 2023; Frazer et al. 2024) to test all possible aa-property combinations across six VPOD datasets. For each dataset, we trained models with twelve ML algorithms and recorded standard performance metrics. Under a null hypothesis of no change between encoding types, we next used Wilcoxon Signed-Rank Tests to compare aa-property and one-hot models based on cross-validation error for the whole dataset (WDS) and on the accuracy of predicting λ_max_ of mutants using models trained only on wild type (WT) data (Gommers et al. 2022; Damian Riina et al. 2023; Frazer et al. 2024). Finally, we built visualization tools to assess the site-wise contributions of specific aa-properties to λ_max_ predictions (Supplemental Methods S3).

### Correcting for phylogenetic signal with Phylogenetically-Weighted Cross-Validation (PW-CV)

To reduce inflated metrics in ML models from phylogenetic non-independence, we developed *‘Phylogenetically-Weighted Cross-Validation’* (PW-CV). PW-CV assign genes to cross-validation folds—subsets of the data used for training or testing in rotation—so that each fold contains phylogenetically dissimilar sequences, to reduce overestimation of accuracy that may occur when closely related sequences appear in both training and testing sets. We created a pairwise phylogenetic distance matrix for all target opsins from a maximum likelihood gene tree estimated with IQ-TREE (Minh et al. 2020). PW-CV initializes folds by selecting the ‘n’ most mutually distant genes, where ‘n’ is the desired number of folds, with remaining genes iteratively added to the fold whose members are most phylogenetically dissimilar, assuming the gene exceeds a minimum distance threshold. This threshold is a user-specified percentile of all pairwise distances in the tree. We defined four ‘relation-handling-methods’ (RHMs: ‘random’, ‘merge’, ‘max_mean’, ‘leave_out’) to manage tips that fell below the threshold for all available folds (see Supplemental Methods S4 for details). Following the complete assignment of all tips to a fold, we then trained ML models using the resulting cross-validation fold assignments.

Using this pipeline, we conducted a large-scale analysis on the wild-type (WT) data subsets from VPOD_1.2, systematically varying the percentile threshold (1–95%), number of folds (5–20), and RHMs to assess impacts on model performance across different ML algorithms and sequence encoding methods. We used the ‘LG+F+R7’ model for constructing the WT opsin gene tree; which was determined by IQ-Tree’s ‘*ModelFinder Plus*’ (MFP+LM) model search parameter. Additionally, we used 1000 ultra-fast bootstrap replicates to assess the statistical support for branches in the resulting tree.

### Optimizing model hyperparameters via grid search

To determine the best-fit hyperparameters for training ML models, we performed comprehensive grid searches (Pedregosa et al. 2011; Liashchynskyi and Liashchynskyi 2019; Baghbanzadeh et al. 2023) for the three algorithms (GBR, XGB, and RF), each applied to six primary VPOD data subsets (WDS, WT, Vert, Invert, WT-Vert, and T1), using both aa-property encoding and one-hot encoding strategies. For aa-property encoding, we first selected the top ten performing aa-property combinations (five by best R², five by lowest MAE) from earlier combinatorial analysis. We recorded grid search-optimized R² and MAE values and their corresponding hyperparameters, ultimately selecting the single best model algorithm and, for aa-property encoded models, the best aa-property combination and ML algorithm for each data subset (see Supplemental Methods S5, Supplementary Tables 3 & 4).

### Mine-N-Match (MNM) Pipeline: Linking genotypes to *in-vivo* phenotypes

We developed the *Mine-N-Match* (MNM) pipeline to address the opsin “data-cliff,” where many more species have opsin sequences than associated λ_max_ measurements. MNM uses *in-vivo* λ_max_ values compiled from prior studies, retrieves opsin sequences from the same species, predicts λ_max_ for those sequences with OPTICS, and expands the amount of sequence data with corresponding phenotypes when predictions match measurements. We first assembled *VPOD_in_vivo_v1.0* (Supplemental Methods S6) from six published compilations and additional literature searches to harmonize *in-vivo* λ_max_ data from microspectrophotometry (Liebman 1972; Bowmaker 1984), electroretinograms (Jacobs et al. 1996; Rocha et al. 2016), and related techniques. As a proof of concept, *VPOD_in_vivo_v1.0* is not comprehensive and additional λ_max_ measures can be added in future updates. Using the species list from *VPOD_in_vivo_v1.0,* we queried the NCBI taxonomy database (Schoch et al. 2020), corrected names with the Global Biodiversity Information Facility (GBIF) taxonomy when necessary (Lane 2003; Edwards 2004), and retrieved opsin coding sequences (nucleotide and protein) from NCBI (Sayers et al. 2025).

We supplemented these from other compiled sources to capture sequences absent from NCBI (Supplemental Methods S6). We ran OPTICS_v1.2 to predict λ_max_ for all candidate sequences and matched each predicted value to the closest *in-vivo* λ_max_ measurement for each species in *VPOD_in_vivo_v1.0*. Matches were filtered to retain only the best match (absolute difference ≤15 nm) and to remove sequences already in VPOD or flagged as unlikely opsins by OPTICS BLASTp output (Supplemental Methods S6). We integrated these genotype-phenotype pairs into VPOD_v1.3, adding new “physiologically inferred” data subsets, each named with an “MNM” suffix in metadata and FASTA files. We trained models on these subsets and compared their performance to models trained on heterologously expressed opsins (see Supplemental Methods S7).

## Results

### OPTICS: An accessible ML tool for predicting opsin spectral sensitivity

The *Opsin Phenotype Tool for Inference of Color Sensitivity* (OPTICS) enables rapid prediction of opsin spectral sensitivity (λ_max_) from protein sequences using machine learning models trained on genotype–phenotype data from the Visual Physiology Opsin Database (VPOD) (Frazer et al. 2024). OPTICS offers multiple pre-trained models tailored to different taxonomic groups, and allowing different sequence encoding strategies, including one-hot and amino acid property (aa-prop) encoding. In addition to λ_max_ prediction, OPTICS provides optional analyses for sequence interpretation, including BLASTp comparisons to training data, site-difference reports standardized to reference sequences, and bootstrap-based confidence estimates that can be visualized as distribution plots (Figure 1). The tool outputs predicted λ_max_ values (mean, median, bounds, and standard deviation) in TSV and Excel formats, along with metadata on the model, encoding method and a parameter log; optional outputs include BLASTp reports and bootstrap plots. OPTICS is freely available via GitHub and as a web application using the Galaxy Project platform (Galaxy Community 2024).

**Figure 1.**
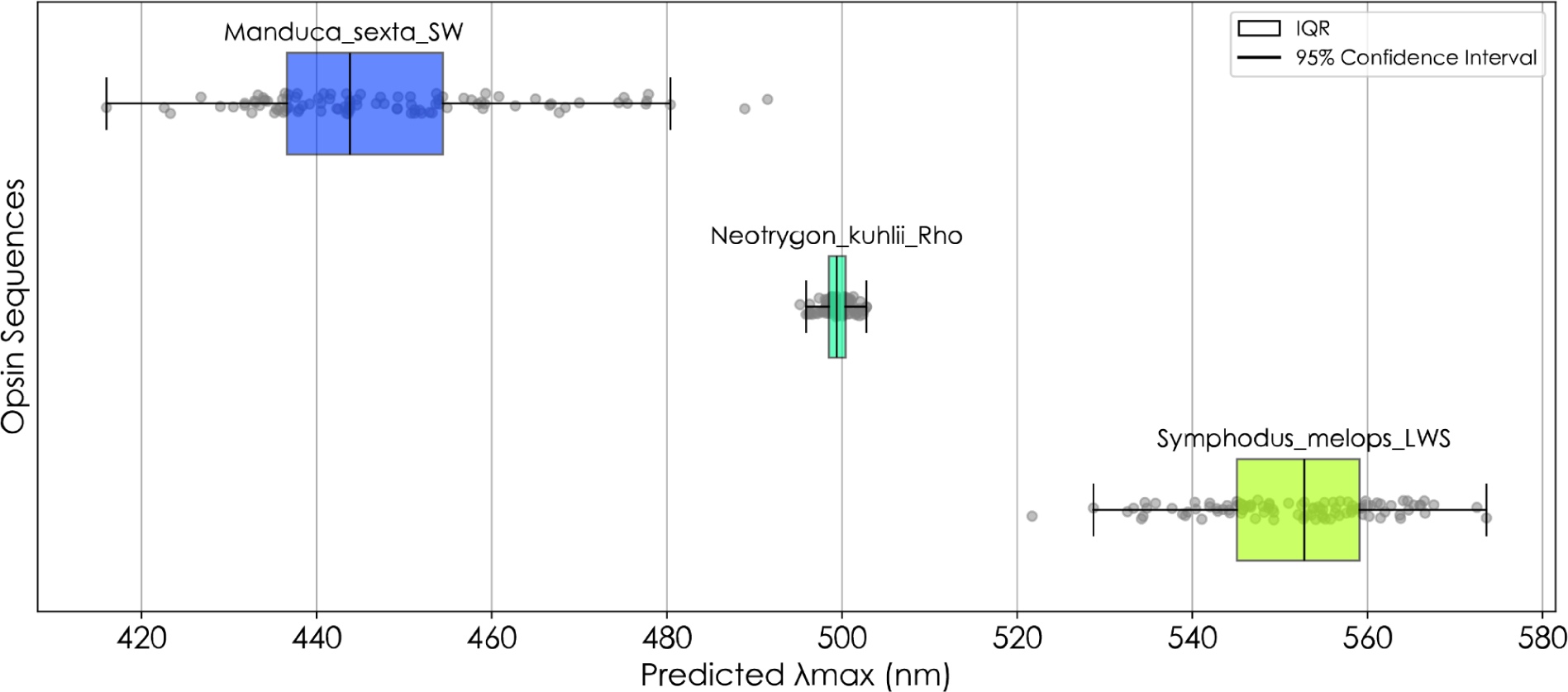
Box-plot distributions of bootstrapped λ_max_ predictions illustrate the contrasting prediction confidence (distribution width) from the WDS bootstrap-ensemble when predicting the λ_max_ for a short-wave (SW) opsin (OX393528.1-9544021-9545976) from the Tobacco Hornworm (*Manduca sexta*)(95% CI Width = 54nm, SD = 14.7), a Rhodopsin (Rho; MN519143.1) from the Bluespotted Stingray (*Neotrygon kuhlii*)(95% CI Width = 6.2nm, SD = 1.6nm), and long-wave sensitive (LWS) opsin (MK983124.1) from the Corkwing Wrasse (Symphodus melops)(95% CI Width = 34.1nm, SD = 10.0nm). Note, the colors of the boxes automatically reflect the median predicted λ_max_ converted to the corresponding hex-code value. This method of uncertainty quantification and visualization formed an integral component for reporting OPTICS prediction uncertainty.

### Bootstrapping provides feedback on prediction stability and confidence

Bootstrapping produced quantitative confidence intervals and allowed qualitative assessment of prediction distributions, as illustrated for the three example opsins (Figure 1). The SW opsin from *M. sexta* had a wide 95% CI (54 nm, SD = 14.7), the Rho from *N. kuhlii* had a narrow CI (6.2 nm, SD = 1.6), and the LWS opsin from *S. melops* was intermediate (34.1 nm, SD = 10.0). In plots, box colors automatically reflect the median predicted λ_max_ converted to the corresponding hex-code value. This quantification and visualization of uncertainty using bootstrapping is an integral feature of OPTICS outputs.

### Encoding with physicochemical properties of amino-acids enhances ML interpretability and predictive power

Across all datasets, models using aa-property encodings achieved higher R² and lower error metrics than one-hot models (Table 1). For example, in WDS, aa-property encoding improved R² from 0.956 to 0.964 and reduced MAE by 18.5% (6.48 nm to 5.28 nm; p = 2.43×10^-14, Wilcoxon test). We also saw some improvement for WT models predicting mutant opsins, with MAE decreasing from 10.0 nm to 9.63 nm (p = 0.046, Wilcoxon test). Optimal aa-property sets differed by dataset, but hydrophilicity and hydrogen bonding properties appeared frequently.

**Table 1.** Performance Metrics Across Opsin Subsets and Top Performing Models Using Amino-Acid Property Encoding.

| Name | Data Subset Version | # Seqs | Top ML Algorithm | Top AA_Prop Combos | <sup>b</sup>R<sup>2</sup> | <sup>a</sup>MAE [nm] | <sup>a</sup>MAPE [%] | <sup>b</sup>MSE | <sup>b</sup>RMSE |
| --- | --- | --- | --- | --- | --- | --- | --- | --- | --- |
| Whole Dataset (WDS) | VPOD_wds_het_1.2 | 1211 | XGB | H1,H2,H3,P2,V,SC<br>T,PKA | 0.964 | 5.49 | 1.23 | 139 | 11.5 |
| Wild-Types (WT) | VPOD_wt_het_1.2 | 364 | GBR | H2,H3,P1,NCI,<br>PKA | 0.939 | 8.01 | 1.73 | 181 | 13.0 |
| Vertebrates (Vert) | VPOD_vert_het_1.2 | 1057 | XGB | H2,H3,NCI,SCT,PK<br>B | 0.978 | 5.13 | 1.14 | 81.5 | 8.91 |
| WT Vertebrates (WT-Vert) | VPOD_wt_vert_het_1.2 | 319 | GBR | H2,P2,V,MASS | 0.979 | 4.73 | 1.01 | 56.2 | 7.14 |
| Invertebrates (Invert) | VPOD_inv_het_1.2 | 155 | GBR | H1,H3 | 0.831 | 13.4 | 2.86 | 515 | 21.2 |
| Whole Dataset MNM (WDS-mnm) | VPOD_wds_het+vivo_1.0 | 1714 | XGB | H2,H3,NCI,<br>MASS | 0.968 | 5.75 | 1.25 | 118 | 10.7 |
| Wild-Types MNM (WT-mnm) | VPOD_wt_het+vivo_1.0 | 867 | GBR | H1,H2,H3,NCI,<br>MASS,PKA | 0.951 | 7.90 | 1.48 | 131 | 11.3 |
| Vertebrates MNM (Vert-mnm) | VPOD_vert_het+vivo_1.0 | 1412 | XGB | H3,P2,SCT,<br>PKA,PKB | 0.983 | 4.36 | 0.97 | 61.8 | 7.71 |
| WT Vertebrates MNM (WT-Vert-mnm) | VPOD_wt_vert_het+vivo_1.0 | 674 | XGB | H2,H3,PKB | 0.982 | 4.52 | 0.96 | 46.1 | 6.72 |
| Invertebrates MNM (Invert-mnm) | VPOD_inv_het+vivo_1.0 | 302 | GBR | H1,P1,SCT | 0.903 | 11.2 | 2.40 | 341 | 17.5 |
| Type-One Opsins (T1) | Karyasuyama_T1_ops | 884 | XGB | H3,P1,PKB | 0.846 | 7.99 | 1.49 | 141 | 11.8 |

**Summary:** Machine learning (ML) models trained on opsin genotype-phenotype data using optimal sets of amino-acid properties are capable of accurately predicting $\lambda_{\text{max}}$ . <sup>a</sup>Mean absolute error (MAE) and mean absolute percent error (MAPE) are in relation to the absolute error $\lambda_{\text{max}}$ predictions and interpreted in the same units of ‘nm’. <sup>b</sup>R<sup>2</sup>, mean square error (MSE) or root mean square error (RMSE) are often interpreted as direct measures of comparing/analyzing model performance and used as training loss terms of the objective function - which measures how well the model fits the training data. One has to often balance between this and the regularization term, which controls the complexity of the model. Thus, a high performance is both simple and predictive; a tradeoff referred to as the ‘bias-variance’ tradeoff.

We also used aa-property encoding to improve biological interpretability. Visualizations of site–property importance (Fig. 2A) showed that amino acid mass at site 43, hydrophobicity at site 94, and charge at site 124 strongly influenced predictions of λ_max_. At site 94 (Fig. 2C), we find higher hydrophobicity to be correlated with lower λ_max_. These property–phenotype relationships were less apparent in standard amino acid identity plots from one-hot encoding (Fig. 2B-D), highlighting the advantage of property-based visualizations for mechanistic insight.

**Figure 2.**
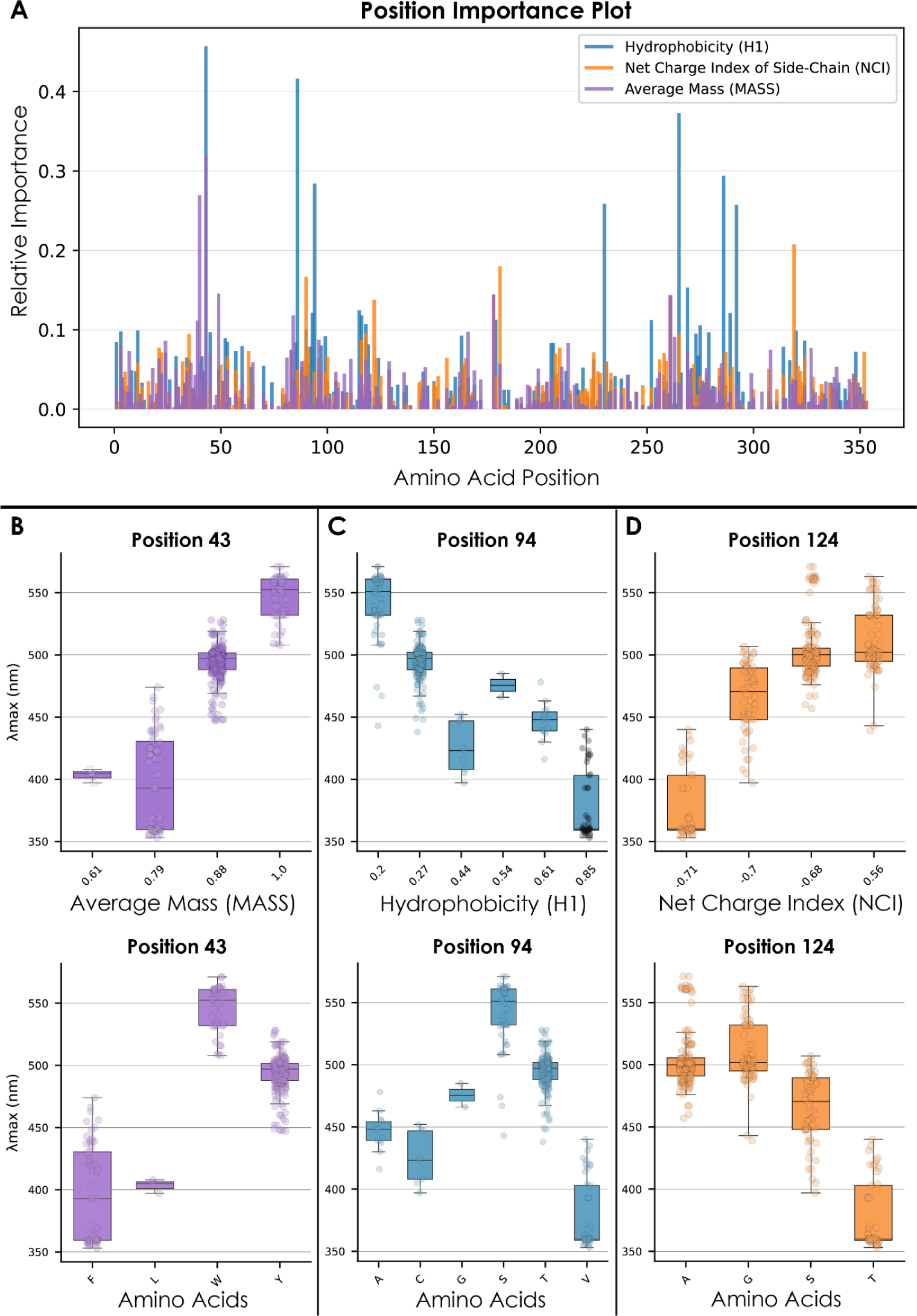
A WT model trained on the amino-acid property (aa-property) combination of, hydrophobicity (H1), net-charge-index (NCI), and mass (MASS), reveals that properties such as amino-acid mass at site 43, hydrophobicity at site 94, and amino-acid charge at site 124 exerted relatively high influence on predictions of λ_max_. **(A)** Bar graph of relative importance by position generated via the mean importance values from the ‘BayesianRidge’, ‘GBR’, ‘XGB’ and ‘RF’ ML regression model trained on the WT-Vertebrate opsin dataset. We interpret positions with higher relative importance as having a larger effect or weight on λ_max_ prediction. Positions 43, 94, and 124 are highlighted with arrows because they are among the highest scoring sites and have all been previously characterized as functionally important to opsin phenotype and function. **(B-D)** Distribution box plots for sites 43, 94 and 124, comparing the relationship between selected aa-property (top) and associated range of λ_max_ values. We also provided corresponding plots for each site when using one-hot encoding (bottom) instead of aa-property encoding; showcasing the difference in immediate biological interpretability. The top distribution box plots provide a biologically interpretable visualization for the relationship between the measure of a physiochemical aa-property and associated range of λ_max_ values at a site of interest. At sites 43 and 124, increasing amino-acid mass and charge values display a positive correlation with true λ_max_ values, while increasing hydrophobicity is negatively correlated with λ_max_ at site 94. This ease of biological interpretation is lost when we simply visualize the relationship between amino-acids and λ_max_. Note, all aa-properties are normalized, either −1-1 or 0-1 for ML training.Thus, these values do not reflect true units. For a more detailed explanation on how position importance scores are calculated for different models, refer to the “*Interpretation*” heading under the methods section of the deepBreaks publication (Baghbanzadeh et al. 2023).

### Phylogenetically-Weighted Cross-Validation (PW-CV) incorporates phylogenetic non-independence in ML performance assessment

*Phylogenetically-Weighted Cross-Validation* (PW-CV) showed a clear link between the stringency of phylogenetic partitioning (percentile threshold) and performance metrics. As thresholds became more stringent and forced greater phylogenetic distance, R² generally declined across most algorithms (Fig. 3A), indicating that standard k-fold cross-validation overestimates generalizability when ignoring phylogenetic relationships. These results were influenced by the specific relation-handing-method for sequences falling below the distance threshold. The *‘leave_out’* method provided the strictest phylogenetic separation within folds, leading to the most conservative evaluation of model performance (Figure 5A/B). But *‘leave_out’* results in a consistent reduction in the size of training datasets, leading to lower absolute performance than*’random’*, or *’max_mean’* methods, which are less strict in their phylogenetic separation (Figure 4B) The *‘merge’* method yielded highly variable outcomes and sometimes underestimated performance due to unbalanced folds. Among training algorithms, Gradient Boosted Regressor (GBR), Extreme Gradient Boosted Regressor (XGB), and Random Forest Regressor (RF) consistently ranked as top performers, whereas Lasso, HuberRegressor, and ExtraTrees performed worst (Fig. 3A), similar to results from Frazer et al. (2024). Changing the number of folds had smaller effects than adjusting thresholds, and comparisons of encoding approaches confirmed that aa-property encoding outperformed one-hot, even under PW-CV (Figure 4C, Supplementary Table 5). Optimization of hyperparameters by grid-search also still improved performance under PW-CV, though absolute values depended on the chosen parameters (Fig. 3D).

**Figure 3.**
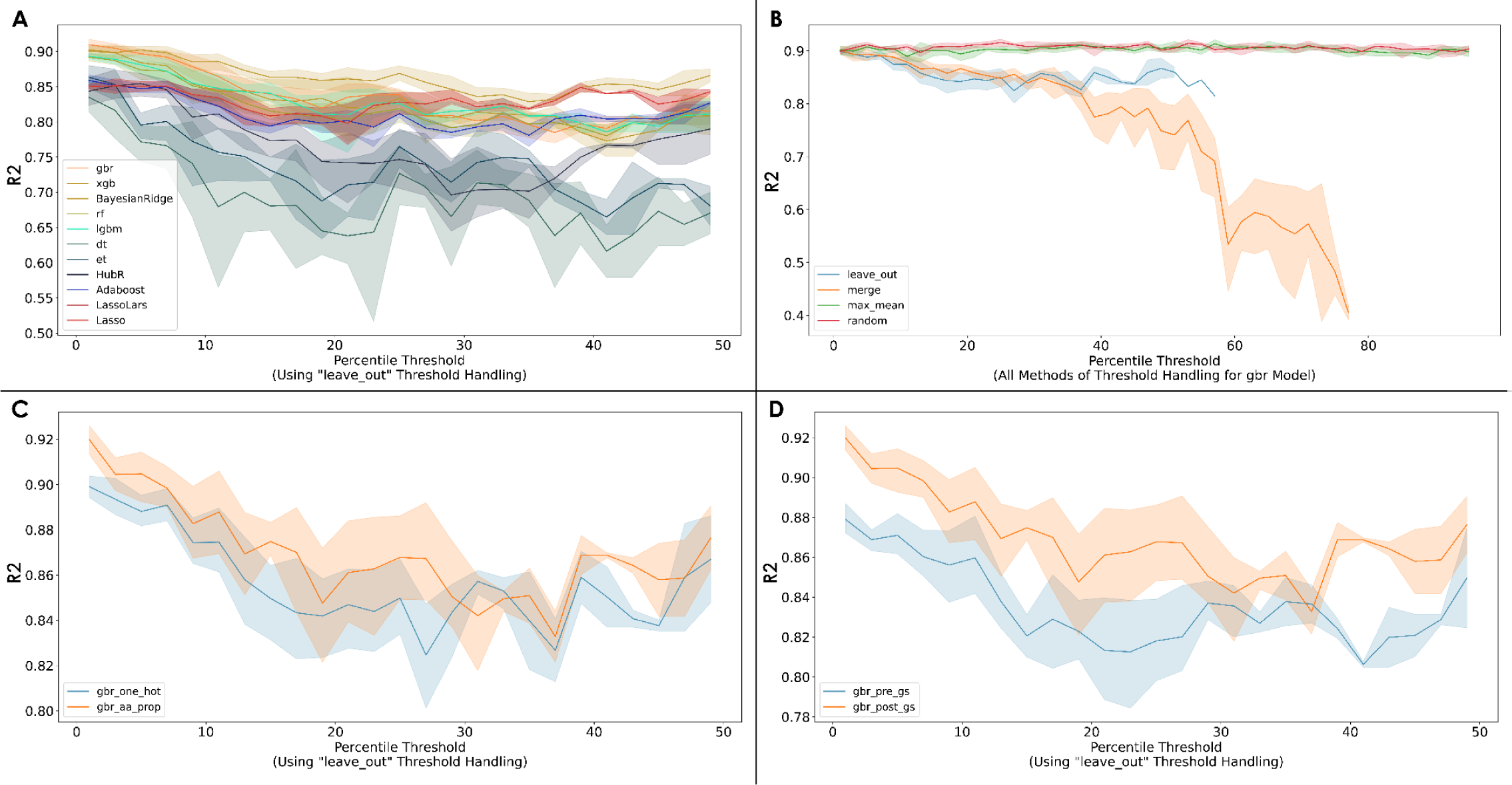
Our analyses using *Phylogenetically-Weighted Cross-Validation* (PW-CV) demonstrated a clear relationship between the stringency of phylogenetic partitioning and model performance metrics. Generally, as the percentile-threshold increased, forcing greater phylogenetic distance between samples within cross-validation folds, we observed a decline in performance metrics such as R² for most algorithms tested. This trend suggests that standard cross-validation approaches, which do not account for phylogeny, may overestimate model generalizability due to the similarity between members of the training and testing splits. **(A-D)** Lighter color ‘error-bars’ represent the spread of model performances (R^2^) relative to the number of cross-validation folds being assigned (5, 8, 10, 12, 15, and 20). **(A)** Line-graph comparison of change in various model R^2^ trends relative to the change in phylogenetic distance threshold. As the threshold increases, the stringency for phylogenetic independence between members of the same cross-validation fold increases. The `‘leave_out’` method shown here discards samples that violate the current phylogenetic distance threshold. (**B)** Line-graph comparing effects of relation-handling-method (RHM) choice on Gradient Boosted Regressor (GBR) model R^2^ relative to the change in phylogenetic distance thresholds. The `‘leave_out’`method stops at a percentile distance threshold of ∼%60 because the resulting datasets created under those thresholds would be too small to train a functional model. For more information on all other RHMs, refer to Supplementary Methods (S3). **(C)** Line-graph comparing the effects of encoding methods used to train ML model (amino-acid properties (orange): gbr_aa_prop, or one-hot (blue): gbr_one_hot) on the change in GBR model R^2^ relative to the change in phylogenetic distance thresholds. The superior performance of models trained with amino acid properties (aa-properties, orange) versus one-hot encoding (blue) is consistent across most phylogenetic distance thresholds. **(D)** Line-graph comparing the effects of ‘pre’ (blue) or ‘post’ (orange) grid-search optimization on the change in GBR model R^2^ relative to the change in phylogenetic distance thresholds. The performance benefit of grid-search hyperparameter optimization (’post’, orange) over default parameters (’pre’, blue) persists even as the stringency of PW-CV increases.

### Hyperparameter optimization via grid search identifies optimal dataset-dependent parameters, enhancing ML predictive power

For models trained with one-hot encoded sequences, our grid search (gs) pipeline identified the specific algorithm (among GBR, XGB, and RF) and its corresponding hyperparameters that yielded the highest R² for each individual dataset. While the best-performing algorithm varied depending on the dataset, the optimization process successfully pinpointed configurations maximizing predictive accuracy based on R². The optimized hyperparameters and the resulting top R² scores for the best-performing algorithm identified for each dataset are detailed in Supplementary Tables 3.

For models trained using aa-property encoded sequences, our grid search pipeline identified the single best-performing combination of aa-properties and ML algorithm hyperparameters for each dataset, based on maximizing the post-grid-search R² value. The specific aa-prop combination selected, the best performing algorithm with its optimized hyperparameters, and the corresponding peak R² values achieved for each dataset are documented in Supplementary Table 4.

### ML can help link genotype data to *in-vivo* measurements of phenotype

*VPOD_in_vivo_v1.0* contains 3810 unique *in-vivo* λ_max_ measurements across 1397 species from nine phyla. From those species, we mined 4523 candidate opsin sequences from 329 species (2308 from NCBI, 2215 from supplementary sources). The MNM pipeline matched predictions from these sequences to physiological λ_max_ values, yielding 695 initial matches. After filtering for best match per species and removing duplicates or poor matches (> 15 nm difference), we retained 555 genotype-phenotype links for use in subsequent model training.

### Harmonizing *in-vivo* and *in-vitro* data improves prediction of spectral sensitivity for undersampled opsin families

We integrated the 555 matches into *VPOD_v1.3*, creating five new data subsets that merge heterologous data and matched data (WDS-mnm, WT-mnm, Vert-mnm, WT-Vert-mnm, Invert-mnm). This addition nearly doubled the number of genotype-phenotype links, as illustrated by the wild-type (WT) (364 to 867) and invertebrate (Invert) (155 to 302) datasets. Models trained on MNM-augmented subsets consistently outperformed those trained on heterologous-only data. For example, a gs-optimized aa-property model on Invert-mnm achieved R² = 0.905, MAE = 10.9 nm, compared to R² = 0.839, MAE = 13.1nm for heterologous-only Invert. WT-mnm models likewise improved (R² = 0.963 vs. 0.939; MAE = 6.70 nm vs. 8.08 nm). Gains were especially strong for understudied invertebrate opsins, suggesting that incorporating inferred *in-vivo* data improves generalization and accuracy (Table 1; Supplementary Tables 2-5). We deployed these new MNM-trained models, plus bootstrap versions, as part of the OPTICS_v1.3 public release on GitHub.

## Discussion

Accurately predicting protein function from sequence remains a central challenge in molecular evolution. We advance this goal using opsin visual pigments by integrating interpretable machine learning, phylogeny-aware evaluation, and systematic data harmonization.

Our contributions include: (1) OPTICS, an accessible and flexible tool to predict λ_max_; (2) encoding based on amino-acid properties that improves accuracy and generalizability over one-hot encoding; (3) bootstrapping and phylogenetically weighted cross-validation (PW-CV) for more realistic estimates of model performance; and (4) the *Mine-N-Match* (MNM) pipeline, which bridges the “data-cliff” between sequence-rich and phenotype-poor datasets. Together, these innovations provide a framework for more accurate genotype-phenotype prediction and make these capabilities broadly available for visual ecology and molecular evolution.

### Democratizing phenotype prediction

Powerful ML models often require specialized expertise, limiting their reach. OPTICS removes this barrier by offering command-line, GUI, and web interfaces that predict λ_max_ directly from amino-acid sequences, while requiring minimal bioinformatics background. The tool packages pre-trained models across data subsets and encoding strategies, automates alignment insertion, and performs bootstrap analysis for confidence intervals. This accessibility enables a wider community, especially visual ecologists and molecular evolutionists, to incorporate predictive modeling into their research.

Despite the broad utility of these ML models, caveats exist. The models predict λ_max_, a central but incomplete descriptor of spectral sensitivity. λ_max_ does not capture activation kinetics or *in-vivo* modifiers like screening pigments (Arikawa and Stavenga 1997), oil-droplets (Das et al. 1999; Hart and Vorobyev 2005; Toomey et al. 2015; Toomey and Corbo 2017), or chromophore shifts (Buczyłko et al. 1996; Terakita 2005; Sekharan and Morokuma 2011). The tool will also return predictions for any sequence, including non-functional or partial opsins, which may yield unreliable results, especially if missing data involve key tuning sites. While OPTICS alerts users when sequence identity to known opsins is low, future versions could incorporate automated filtering of non-opsins and pseudogenes. As with any ML tool, predictions for highly divergent sequences require caution, such that bootstrap intervals will provide critical feedback for assessing reliability.

### Advancing ML genotype–phenotype links: Confidence, phylogeny-aware generalization, and interpretability

Reliable genotype–phenotype prediction from biological sequences requires three pillars: robust confidence estimates, evaluation methods that account for evolutionary relatedness, and biologically grounded interpretability. Confidence intervals provide important context for the reliability of predictions. Herein, bootstrapping demonstrated substantial variance in confidence across different opsins (Figure 1); reflecting differences in how well-sampled are different regions of genotype-phenotype space. Without confidence intervals, predictions from sparsely sampled paralogs could appear as certain as those for conserved, highly sampled genes, like vertebrate rhodopsins.

Also important to reliability is evaluating models in a way that accounts for evolutionary non-independence. Standard cross-validation often inflates performance because closely related sequences can appear in both training and test sets (Roberts et al. 2017). Our phylogenetically weighted cross-validation (PW-CV) examines this bias by incorporating sequence-based phylogenetic distances into the partitioning process. As phylogenetic separation increases, performance metrics generally decline, yielding more conservative estimates of generalization to novel sequences. Importantly, there is no universally optimal phylogenetic threshold because higher thresholds exclude more closely related sequences, increasing conservatism, but eventually reduce the dataset to the point where accuracy declines because less training data are available. This makes the selection of a single “best” threshold inherently arbitrary. Instead of a single threshold, researchers could report performance across a range of thresholds, or examine overall trends. For example, large early drops in performance at low thresholds may indicate a phylogenetically imbalanced dataset in which many closely related genes are rapidly removed.

We found model interpretability to benefit from encoding strategies that reflect amino acid properties. One-hot encoding, while common, treats each amino acid as an unrelated category and fails to capture similarities based on shared physicochemical properties. By contrast, our amino-acid property encoding improves both predictive accuracy and generalization to mutant sequences (Table 1, Supplementary Table 2; Figure 3A/B), likely because it allows the model to exploit biologically relevant relationships, even for amino acids absent from the training set, and because amino acid properties may contain more information about changes to opsin tuning sites. Visualizations of site-wise importance from these models link predictive patterns to known or candidate spectral tuning mechanisms. For example, our models identified hydrophobicity at site 94, a position in the binding pocket near the counterion at site 113 and the Schiff base linkage at site 296, as negatively correlated with λ_max_, consistent with the role of hydrophobic interactions in stabilizing the chromophore binding pocket (Takahashi and Ebrey 2003; Chinen et al. 2005; Tsukamoto et al. 2017). Similarly, charge at site 124, another known tuning site (Zheng et al. 2015; Castiglione and Chang 2018), showed a positive correlation with λ_max_. Such property–phenotype relationships provide direct, mechanistic hypotheses for experimental testing, and future work could also integrate epistasis or multi-property effects to further enhance interpretability.

### Bridging the data-cliff with Mine-N-Match

A persistent challenge in functional genomics is the data cliff, the steep disparity between the abundance of sequence data and the relative scarcity of experimentally characterized phenotypes. This imbalance limits our ability to fully use genomic information for predictive modeling of phenotypes. Opsins exemplify this challenge: while *in-vitro* measurements of heterologously expressed sequences provide precise and controlled functional data, they are demanding technically, especially for invertebrate r-opsins, and thus limited in number. By contrast, *in-vivo* measurements of spectral sensitivity (e.g., microspectrophotometry, electroretinograms) are more numerous across species but are rarely linked to the exact opsin sequences responsible.

The *Mine-N-Match* (MNM) pipeline addresses this gap by using predictions from machine learning as an informed bridge between unlinked *in-vivo* phenotypes and available sequences. MNM automates the retrieval of candidate opsin sequences for species with known *in-vivo* λ_max_, predicts λ_max_ for each candidate using OPTICS, and assigns the most likely sequence to a phenotype. This systematic approach accelerates what has traditionally been a slow, manual process of linking data scattered across public repositories and the literature. By combining automated NCBI queries, taxonomic reconciliation, and ML-based matching, MNM offers an efficient workflow for harmonizing heterogeneous datasets.

Integrating MNM-derived genotype–phenotype pairs into VPOD_v1.3 substantially expanded data coverage, doubling the number of wild-type opsins with linked phenotypes and nearly doubling invertebrate coverage—which are very undersampled in heterologous studies. Models trained on these expanded datasets showed considerable gains, especially for invertebrate r-opsins, where predictive accuracy (R²) and error rates improved relative to models trained solely on *in-vitro* data (Table 1). These results demonstrate that leveraging *in-vivo* data—when genotype links are inferred by robust ML methods—can meaningfully improve prediction accuracy and generalization in data-sparse regions of sequence space.

Although developed for opsins, the MNM strategy could be more broadly applicable to other molecular systems where phenotypes are measured independently of genotypes—such as enzymes with kinetic parameters from biochemical assays but no linked gene sequence, or receptors with pharmacological profiles but unsequenced sources. By pairing systematic sequence mining with predictive modeling, analogous pipelines could help bridge similar data-cliffs across diverse areas of biology, accelerating the integration of legacy phenotypic data into modern sequence-based frameworks.

While the results of the MNM pipeline are promising, it is important to note that success relies on the accuracy of OPTICS predictions to correctly link genotypes to *in-vivo* phenotypes. While we filter matches based on prediction confidence and proximity, incorrect links are still possible, especially for species with multiple opsins having similar predicted λ_max_ values close to an *in-vivo* measurement. The accuracy of the underlying *in-vivo* data in *VPOD_in_vivo_v1.0* and the taxonomic information in NCBI also influences the quality and abundance of these links.

Future work could involve testing the validity of matches by expressing opsins heterologously and comparing the *in-vitro* λ_max_ to the *in-vivo* measurement it was linked to by MNM. Moreover, there were a rather large number of λ_max_ values from *VPOD_in_vivo_1.0* that were not linked to any opsin sequences. These could be a good target for future empirical work.

### Implications & Future Directions

The methods and tools developed here have immediate relevance for visual ecology, sensory biology, molecular evolution, and protein engineering. OPTICS and the expanded *VPOD_v1.3* database can accelerate characterization of visual systems across taxa, enabling large-scale comparative studies and deeper insight into how visual sensitivities adapt to diverse ecological niches. The principles behind PW-CV and amino-acid property encoding are broadly transferable to predictive modeling of other protein functions.

Looking forward, this framework lays the ground for more ambitious explorations of genotype-phenotype landscapes. We envision extending this work into an integrated evolution simulation pipeline (Wittmann et al. 2021; Howard et al. 2025) designed to probe historical contingency and adaptive trajectories in molecular evolution. By simulating evolutionary paths under different varying selective pressures, this pipeline could reveal how different genetic starting points might constrain or expand accessible phenotypes (Upton et al. 2021; Murphy and Westerman 2022), identify repeatable mutational paths to the same phenotype, and show how these paths depend on genotype and selection. Such simulations could help illuminate the predictability of evolution, the role of epistasis in shaping adaptive landscapes, and the molecular underlying visual diversity, further enhancing the role of opsins as a model for broader principles of the molecular evolution of functional diversity.

## Data Availability

All methods, results, and analyses from this study and our previous publication are available in the GitHub repository *The Visual Physiology Opsin Database (VPOD)*: https://github.com/VisualPhysiologyDB/visual-physiology-opsin-db. *Opsin Phenotype Tool for Inference of Color Sensitivity* (OPTICS), is available at: https://github.com/VisualPhysiologyDB/optics. All data and code are released under the GNU General Public License v3.0 (Open Source Initiative–approved). These resources also support applications using the *deepBreaks* ML framework (Baghbanzadeh et al. 2023) and the VPOD database for opsin research (Frazer et al. 2024).

