## Supplemental Material for "Accessible and Robust Machine Learning Approaches to Improve the Opsin Genotype-Phenotype Map"

### Supplementary Material

**Supplementary Table 1.** Table of Scaled Amino Acids and their corresponding Physiochemical Properties

| | H1 | H2 | H3 | V | P1 | P2 | SASA | NCI | MASS | PKA | PKB | [a]Side Chain Type |
|---|---|---|---|---|---|---|---|---|---|---|---|---|
| A | 0.61 | -0.09 | 0.50 | 0.19 | 0.62 | 0.11 | 0.44 | -0.68 | 0.38 | 2.34 | 9.69 | 3 |
| C | 0.44 | -0.25 | 0.50 | 0.31 | 0.42 | 0.31 | 0.55 | -1.00 | 0.55 | 1.96 | 10.28 | 1 |
| D | -0.17 | 1.00 | 1.00 | 0.27 | 1.00 | 0.26 | 0.60 | -0.91 | 0.62 | 1.88 | 9.60 | 6 |
| E | -0.08 | 1.00 | 1.00 | 0.43 | 0.95 | 0.37 | 0.70 | -0.69 | 0.69 | 2.19 | 9.67 | 6 |
| F | 0.90 | -0.72 | 0.50 | 0.79 | 0.40 | 0.71 | 0.84 | -0.46 | 0.79 | 1.83 | 9.13 | 2 |
| G | 0.54 | 0.06 | 0.50 | 0.00 | 0.69 | 0.00 | 0.33 | 0.56 | 0.31 | 2.34 | 9.60 | 5 |
| H | 0.09 | -0.09 | 1.00 | 0.54 | 0.80 | 0.56 | 0.76 | -0.81 | 0.74 | 1.82 | 9.17 | 6 |
| I | 1.00 | -0.50 | 0.50 | 0.64 | 0.40 | 0.45 | 0.68 | -0.58 | 0.61 | 2.36 | 9.60 | 3 |
| K | -0.47 | 1.00 | 0.50 | 0.69 | 0.87 | 0.54 | 0.85 | -0.61 | 0.69 | 2.18 | 8.95 | 6 |
| L | 0.84 | -0.50 | 0.50 | 0.64 | 0.38 | 0.45 | 0.73 | -0.36 | 0.61 | 2.36 | 9.60 | 3 |
| M | 0.62 | -0.34 | 0.50 | 0.65 | 0.44 | 0.54 | 0.76 | -0.72 | 0.70 | 2.28 | 9.21 | 3 |
| N | -0.10 | 0.69 | 1.00 | 0.40 | 0.89 | 0.33 | 0.62 | -0.70 | 0.61 | 2.02 | 8.80 | 4 |
| P | 0.36 | 0.06 | 0.50 | 0.29 | 0.62 | 0.32 | 0.55 | 1.00 | 0.52 | 1.99 | 10.60 | 5 |
| Q | -0.14 | 0.13 | 1.00 | 0.55 | 0.81 | 0.44 | 0.73 | -0.38 | 0.69 | 2.17 | 9.13 | 4 |
| R | -1.00 | 1.00 | 1.00 | 0.72 | 0.81 | 0.44 | 0.73 | -0.38 | 0.84 | 2.17 | 9.04 | 6 |
| S | 0.20 | 0.16 | 1.00 | 0.20 | 0.71 | 0.15 | 0.49 | -0.70 | 0.47 | 2.21 | 9.15 | 4 |
| T | 0.27 | -0.06 | 1.00 | 0.35 | 0.66 | 0.26 | 0.57 | -0.71 | 0.54 | 2.09 | 9.10 | 4 |
| V | 0.85 | -0.41 | 0.50 | 0.49 | 0.45 | 0.34 | 0.62 | -0.32 | 0.53 | 2.32 | 9.62 | 3 |
| W | 0.71 | -1.00 | 0.75 | 1.00 | 0.42 | 1.00 | 1.00 | -0.46 | 1.00 | 2.83 | 9.39 | 2 |
| Y | 0.43 | -0.66 | 0.75 | 0.81 | 0.48 | 0.73 | 0.89 | -0.56 | 0.88 | 2.20 | 9.11 | 2 |

[a]Side Chain Type (SCT) is ranked by reactivity: 1=Cleft-Cysteine, 2=Cleft-Aromatic, 3=Cleft-Aliphatic. 4=Cleft-polar uncharged, 5=Cleft-Pro & Gly, 6=All else

We selected twelve physicochemical properties of amino acids (aa-properties): hydrophobicity (H1), hydrophilicity (H2), hydrogen bonding (H3), polarity (P1), polarizability (P2), volume (V), net charge index of side chains (NCI), average mass of the amino-acid(MASS), solvent-accessible surface area (SASA), acid-dissociation constant (PkA), base-dissociation constant (PkB), and side-chain type (SCT). All properties, except for SCT, were normalized (0-1 or -1-1).

**Supplementary Table 2.** Performance Metrics Across Opsin Subsets and Top Performing Models Using One-Hot Encoding.

| Name | Data Subset Version | # Seqs | Top ML Algorithm | [b]$R^2$ | [a]MAE [nm] | [a]MAPE [%] | [b]MSE | [b]RMSE |
|---|---|---|---|---|---|---|---|---|
| Whole Dataset (WDS) | *VPOD_wds_het_1.2* | 1211 | GBR | 0.956 | 7.31 | 1.69 | 157 | 12.4 |
| Wild-Types (WT) | *VPOD_wt_het_1.2* | 364 | GBR | 0.912 | 9.43 | 2.04 | 257 | 15.4 |
| Vertebrates (Vert) | *VPOD_vert_het_1.2* | 1057 | XGB | 0.971 | 6.02 | 1.26 | 110 | 10.3 |
| WT Vertebrates (WT-Vert) | *VPOD_wt_vert_het_1.2* | 319 | GBR | 0.973 | 5.22 | 1.14 | 72.6 | 8.08 |
| Invertebrates (Invert) | *VPOD_inv_het_1.2* | 155 | BR | 0.808 | 15.0 | 3.27 | 616 | 23.4 |
| Whole Dataset MNM (WDS-mnm) | *VPOD_wds_het+vivo_1.0* | 1714 | XGB | 0.960 | 6.32 | 1.40 | 148 | 12.0 |
| Wild-Types MNM (WT-mnm) | *VPOD_wt_het+vivo_1.0* | 867 | GBR | 0.946 | 8.34 | 1.76 | 164 | 12.7 |
| Vertebrates MNM (Vert-mnm) | *VPOD_vert_het+vivo_1.0* | 1412 | XGB | 0.981 | 4.79 | 1.07 | 71.4 | 8.27 |
| WT Vertebrates MNM (WT-Vert-mnm) | *VPOD_wt_vert_het+vivo_1.0* | 674 | XGB | 0.979 | 4.67 | 0.99 | 53.0 | 6.95 |
| Invertebrates MNM (Invert-mnm) | *VPOD_inv_het+vivo_1.0* | 302 | GBR | 0.878 | 12.8 | 2.71 | 431 | 19.5 |
| Type-One Opsins (T1) | *Karyasuyama_T1_ops* | 884 | XGB | 0.803 | 9.05 | 1.69 | 180 | 13.3 |

[a]*Mean absolute error* (MAE) and *mean absolute percent error* (MAPE) are in relation to the absolute error $\lambda_{max}$ predictions and interpreted in the same units of 'nm'. [b]$R^2$, *mean square error* (MSE) or *root mean square error* (RMSE) are often interpreted as direct measures of comparing/analyzing model performance and used as training loss terms of the objective function - which measures how well the model fits the training data. One has to often balance between this and the regularization term, which controls the complexity of the model. Thus, a high performance is both simple and predictive; a tradeoff referred to as the '*bias-variance'* tradeoff.

**Supplementary Table 3.** Grid-Search Optimized Parameters Across Opsin Subsets and Top Performing Models Using One-Hot Encoding.

| Name | Top ML Algorithm | gs-$R^2$ | gs-MAE | gs-MSE | gs-Optimized Model Params |
|---|---|---|---|---|---|
| Whole Dataset (WDS) | GBR | 0.961 | 6.48 | 155 | {'gbr__learning_rate': 0.2, 'gbr__max_depth': 3, 'gbr__max_features': 'sqrt', 'gbr__n_estimators': 500} |
| Wild-Types (WT) | XGB | 0.921 | 8.69 | 232 | {'xgb__gamma': 1.0, 'xgb__learning_rate': 0.1, 'xgb__max_depth': 3, 'xgb__n_estimators': 300, 'xgb__reg_alpha': 0, 'xgb__reg_lambda': 0} |
| Vertebrates (Vert) | XGB | 0.976 | 6.05 | 92.4 | {'xgb__gamma': 0, 'xgb__learning_rate': 0.1, 'xgb__max_depth': 3, 'xgb__n_estimators': 200, 'xgb__reg_alpha': 1.0, 'xgb__reg_lambda': 1.0} |
| WT Vertebrates (WT-Vert) | GBR | 0.975 | 5.04 | 67.4 | {'gbr__learning_rate': 0.2, 'gbr__max_depth': 3, 'gbr__max_features': None, 'gbr__n_estimators': 200} |
| Invertebrates (Invert) | BR | 0.808 | 15.0 | 616 | {'BayesianRidge__alpha_1': 0.01, 'BayesianRidge__alpha_2': 1e-06, 'BayesianRidge__compute_score': True, 'BayesianRidge__fit_intercept': True, 'BayesianRidge__lambda_1': 1e-06, 'BayesianRidge__lambda_2': 0.01} |
| Whole Dataset MNM (WDS-mnm) | XGB | 0.967 | 5.96 | 121 | {'xgb__colsample_bytree': 0.8, 'xgb__gamma': 0, 'xgb__learning_rate': 0.1, 'xgb__max_depth': 5, 'xgb__n_estimators': 200, 'xgb__reg_alpha': 1.0, 'xgb__reg_lambda': 0.1, 'xgb__subsample': 1.0} |
| Wild-Types MNM (WT-mnm) | GBR | 0.960 | 6.93 | 122 | {'gbr__learning_rate': 0.1, 'gbr__max_depth': 5, 'gbr__max_features': 'sqrt', 'gbr__n_estimators': 500} |
| Vertebrates MNM (Vert-mnm) | XGB | 0.982 | 4.63 | 64.6 | {'xgb__colsample_bytree': 0.8, 'xgb__gamma': 1.0, 'xgb__learning_rate': 0.2, 'xgb__max_depth': 5, 'xgb__n_estimators': 100, 'xgb__reg_alpha': 0.1, 'xgb__reg_lambda': 1.0, 'xgb__subsample': 1.0} |
| WT Vertebrates MNM (WT-Vert-mnm) | XGB | 0.983 | 4.41 | 43.7 | {'xgb__colsample_bytree': 1.0, 'xgb__gamma': 0.1, 'xgb__learning_rate': 0.2, 'xgb__max_depth': 5, 'xgb__n_estimators': 100, 'xgb__reg_alpha': 0.1, 'xgb__reg_lambda': 1.0, 'xgb__subsample': 1.0} |
| Invertebrates MNM (Invert-mnm) | GBR | 0.898 | 11.5 | 365 | {'gbr__learning_rate': 0.01, 'gbr__max_depth': 5, 'gbr__max_features': 'sqrt', 'gbr__n_estimators': 500} |
| Type-One Opsins (T1) | XGB | 0.820 | 9.03 | 166 | {'xgb__colsample_bytree': 0.8, 'xgb__gamma': 0, 'xgb__learning_rate': 0.2, 'xgb__max_depth': 3, 'xgb__n_estimators': 200, 'xgb__reg_alpha': 1.0, 'xgb__reg_lambda': 0, 'xgb__subsample': 0.8} |

**Supplementary Table 4.** Grid-Search Optimized Parameters Across Opsin Subsets and Top Performing Models Using Amino-Acid Property Encoding.

| Name | Top ML Algorithm | gs-$R^2$ | gs-MAE | gs-MSE | gs-Optimized Model Params |
|---|---|---|---|---|---|
| Whole Dataset (WDS) | XGB | 0.969 | 5.28 | 117 | {'xgb__colsample_bytree': 0.8, 'xgb__gamma': 0, 'xgb__learning_rate': 0.2, 'xgb__max_depth': 5, 'xgb__n_estimators': 100, 'xgb__reg_alpha': 0.1, 'xgb__reg_lambda': 0.1, 'xgb__subsample': 1.0} |
| Wild-Types (WT) | GBR | 0.939 | 8.08 | 180 | {'gbr__learning_rate': 0.1, 'gbr__max_depth': 3, 'gbr__max_features': 'log2', 'gbr__n_estimators': 500} |
| Vertebrates (Vert) | XGB | 0.981 | 4.95 | 72.0 | {'xgb__colsample_bytree': 1.0, 'xgb__gamma': 0, 'xgb__learning_rate': 0.1, 'xgb__max_depth': 5, 'xgb__n_estimators': 200, 'xgb__reg_alpha': 0.1, 'xgb__reg_lambda': 0.1, 'xgb__subsample': 1.0} |
| WT Vertebrates (WT-Vert) | GBR | 0.980 | 4.60 | 54.7 | {'gbr__learning_rate': 0.1, 'gbr__max_depth': 3, 'gbr__max_features': None, 'gbr__n_estimators': 300} |
| Invertebrates (Invert) | GBR | 0.839 | 13.1 | 498 | {'gbr__learning_rate': 0.01, 'gbr__max_depth': 3, 'gbr__max_features': None, 'gbr__n_estimators': 500} |
| Whole Dataset MNM (WDS-mnm) | XGB | 0.970 | 5.64 | 109 | {'xgb__colsample_bytree': 0.8, 'xgb__gamma': 1.0, 'xgb__learning_rate': 0.1, 'xgb__max_depth': 5, 'xgb__n_estimators': 300, 'xgb__reg_alpha': 0, 'xgb__reg_lambda': 0, 'xgb__subsample': 1.0} |
| Wild-Types MNM (WT-mnm) | GBR | 0.963 | 6.70 | 112 | {'gbr__learning_rate': 0.1, 'gbr__max_depth': 3, 'gbr__max_features': 'sqrt', 'gbr__n_estimators': 800} |
| Vertebrates MNM (Vert-mnm) | XGB | 0.986 | 4.16 | 53.4 | {'xgb__colsample_bytree': 0.8, 'xgb__gamma': 1.0, 'xgb__learning_rate': 0.1, 'xgb__max_depth': 5, 'xgb__n_estimators': 300, 'xgb__reg_alpha': 0, 'xgb__reg_lambda': 0.1, 'xgb__subsample': 0.8} |
| WT Vertebrates MNM (WT-Vert-mnm) | XGB | 0.986 | 4.09 | 35.2 | {'xgb__colsample_bytree': 0.8, 'xgb__gamma': 0, 'xgb__learning_rate': 0.1, 'xgb__max_depth': 5, 'xgb__n_estimators': 200, 'xgb__reg_alpha': 1.0, 'xgb__reg_lambda': 1.0, 'xgb__subsample': 0.8} |
| Invertebrates MNM (Invert-mnm) | GBR | 0.905 | 10.9 | 337 | {'gbr__learning_rate': 0.01, 'gbr__max_depth': 5, 'gbr__max_features': 'sqrt', 'gbr__n_estimators': 500} |
| Type-One Opsins (T1) | XGB | 0.869 | 7.55 | 121 | {'xgb__colsample_bytree': 0.8, 'xgb__gamma': 0.1, 'xgb__learning_rate': 0.1, 'xgb__max_depth': 5, 'xgb__n_estimators': 300, 'xgb__reg_alpha': 1.0, 'xgb__reg_lambda': 1.0, 'xgb__subsample': 0.8} |

**Supplementary Table 5.** *Phylogenetically-Weighted Cross-Validation* Performance Metrics For Grid-Search Optimized Models Trained on Both One-Hot and AA-Property Encoded Sequence Data

| Name | Encoding Method | Top ML Algorithm | Grid-Search Optimized? | $R^2$ 5% DPT | $R^2$ 11% DPT | $R^2$ 21% DPT | MAE 5% DPT | MAE 11% DPT | MAE 21% DPT |
|---|---|---|---|---|---|---|---|---|---|
| Wild-Types (WT) | One-Hot | XGB | N | 0.884 | 0.844 | 0.820 | 11.1 | 13.6 | 15.9 |
| | | | Y | 0.900 | 0.864 | 0.844 | 10.8 | 13.0 | 14.9 |
| | AA-Properties | GBR | N | 0.871 | 0.860 | 0.812 | 11.7 | 13.6 | 16.0 |
| | | | Y | 0.905 | 0.888 | 0.861 | 10.2 | 12.0 | 13.5 |
| Wild-Types MNM (WT-mnm) | One-Hot | XGB | N | 0.933 | 0.916 | 0.846 | 9.84 | 10.8 | 12.2 |
| | | | Y | 0.948 | 0.924 | 0.864 | 8.41 | 10.0 | 11.2 |
| | AA-Properties | GBR | N | 0.941 | 0.921 | 0.858 | 9.32 | 10.5 | 11.4 |
| | | | Y | 0.950 | 0.930 | 0.885` | 8.19 | 9.58 | 10.3 |

**All $R^2$ and MAE values are calculated as the average of the values obtained from running PW-CV using 5, 8, 10, 12, 15, and 20 folds respectively.*

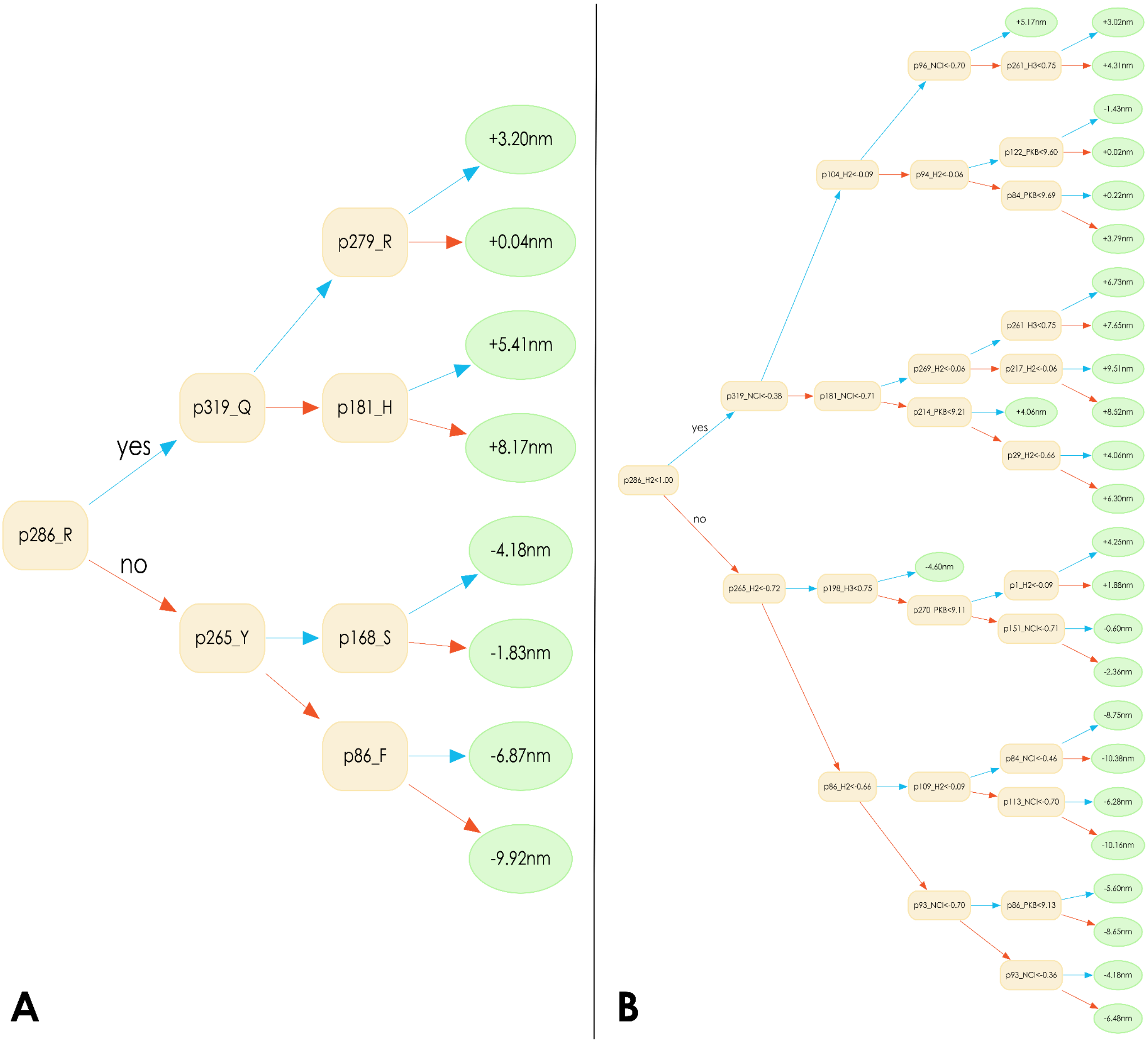


**Supplementary Figure 1. (A)** Decision-tree visualization from a singular decision tree of XGB model trained using One-Hot encoding on the WT dataset from VPOD_1.2 **(B)** Decision-tree visualization from a singular decision tree of XGB model trained using Amino-Acid Property encoding on the WT dataset from VPOD_1.2

### Supplemental Methods

#### S1. OPTICS Implementation and Availability

*OPTICS_v1.2* was implemented in Python (v3.11.9) and relies on libraries including *Scikit-learn* and *BioPython* (see *requirements.txt* on the OPTICS GitHub). External dependencies are NCBI BLAST+ (Camacho et al. 2009) and MAFFT (Katoh and Standley 2013); MAFFT is bundled for Windows. Available models in *OPTICS_v1.2* (and their command-line references) include those trained on WDS (*whole-dataset*), WT (*wildtype*), Vert (*vertebrate*), WT-Vert (*wildtype-vert*), Invert (*invertebrate*), and T1 (*type-one*) datasets. OPTICS uses *Joblib* and *Multiprocessing* for parallelized predictions. Output directories are standardized as *optics_on_{output_dir}_{timestamp}*. Source code is available at GitHub (https://github.com/VisualPhysiologyDB/optics) under GPL v3. A web version is hosted on a Galaxy Platform server (Galaxy Community 2024)[http://galaxy-dev.cnsi.ucsb.edu:8080/?tool_id=optics_1&version=latest]. OPTICS is an ongoing project and may accumulate more features than originally described in this publication.

#### S2. Bootstrapping Pipeline Details

Bootstrap ensembles were generated for WDS, WT, Vert, Invert, and WT-Vert VPOD data subsets. Individual bootstrapped models were saved as PKL files (e.g., *gbr_69.pkl*) in dataset-specific folders (e.g., *wds_bootstrap*). Scripts are in *scripts_n_notebooks/vpod_ML_workflows/bootstrap_model_gen* on the VPOD GitHub repository.

#### S3. Physicochemical amino-acidProperties

We selected twelve physicochemical properties of amino acids (aa-properties): hydrophobicity (H1), hydrophilicity (H2), hydrogen bonding (H3), polarity (P1), polarizability (P2), volume (V), net charge index of side chains (NCI), average mass of the amino-acid(MASS), solvent-accessible surface area (SASA), acid-dissociation constant (PkA), base-dissociation constant (PkB), and side-chain type (SCT). Ten of these were used in previous studies (Li et al. 2016; Inoue et al. 2021), PkA and PkB were sourced from *MilliporeSigma* reference charts, and SCT was a categorical property we defined. All properties, except for SCT, were normalized (0-1 or -1-1) in-line with a similar implementation of this methodology (Li et al. 2016). The *AminoAcidPropertyEncoder* function was integrated into the *deepBreaks* pipeline (Baghbanzadeh et al. 2023; Frazer et al. 2024). Helper functions *aaprop_importance_from_pipe* and *dp_aa_prop_plot* were written for visualizing site-wise property importance and relationships with $\lambda_{max}$, respectively. These functions are available in *preprocessing.py* and *visualization.py* scripts within the *deepBreaks* folder in our VPOD GitHub repository (*scripts-n-notebooks/vpod_ML_workflows/vpod_scripts*). The combinatorial aa-property encoding experiment workflow, *aa_prop_combos.py*, can be found under *scripts-n-notebooks/vpod_ML_workflows/subtests* in VPOD.

#### S4. Phylogenetically-Weighted Cross-Validation (PW-CV) Details.

The four relation-handling-methods (RHMs) were:

- ***'random'***: Assigns a tip violating the distance-threshold to a random fold.
- ***'merge'***: Assigns the tip to the fold containing the member to which it is most closely related (smallest phylogenetic distance).
- ***'max_mean'***: Assigns the tip to the fold with the greatest mean phylogenetic distance from it.
- ***'leave_out'***: Assigns a fold identity of '-1' to the tip, and these tips are later excluded from training.

All scripts for PW-CV are in *scripts_n_notebooks/vpod_ML_workflows/subtests/phylo_weighted_cv* and results are in *result_files/phylo_weighted_cv* on the VPOD GitHub repository.

#### S5. Hyperparameter Optimization (Grid Search) Details

The full set of ML algorithm-specific hyperparameters explored during grid search can be found in the *get_exp_params* module in *deepBreaks/utils_alt.py* on the VPOD GitHub repository. For one-hot encoded models, results are in Supplementary Table 3. For aa-property encoded models, the top five combinations by $R^2$ and top five by MAE from the combinatorial analysis were subjected to grid search; final results are in Supplementary Table 4. Scripts (*vpod_one_hot_grid_search_iter.py* and *vpod_aa_prop_grid_search_iter.py*) are in *scripts_n_notebooks/vpod_ML_workflows/subtests/grid_search* in VPOD.

#### S6. Mine-N-Match (MNM) Pipeline Details

***VPOD_in_vivo_v1.0***: The individual components of *VPOD_in_vivo_v1.0* were sourced from the *Longcore (Longcore 2023)*, *Murphy_Westerman (Murphy and Westerman 2022)*, *Caves_Fish (Schweikert et al. 2019), Porter_1 (Porter 2005), Porter_2 (Porter et al. 2007)*, and *J_Kooi (van der Kooi et al. 2021)* publications; in combination with our custom curated dataset. Our custom curated dataset of *in-vivo* $\lambda_{max}$ data was created following similar methods in Frazer et al. (2024), cataloging species, phylum, $\lambda_{max}$ , error, cell type, chromophore, life-stage, and literature source (linked to VPOD *litsearch* table) in *VPOD_in_vivo_data.csv.* However, one should note that this custom data-set was not originally compiled for the purpose of this study; rather we repurposed this data in an *ad-hoc* manner. As such, we do not label this dataset as an independent element of VPOD which can be versioned in the same way as our heterologous datasets. We used the *merge_accessory_dbs* function in *mine_n_match_functions.py* (located under the directory *scripts_n_notebooks/vpod_ML_workflows/mine_n_match/mnm_scripts* in VPOD) to harmonize these data collections into a single compendium. We deemed this resulting compendium of *in-vivo* $\lambda_{max}$ data, *VPOD_in_vivo_v1.0* (which can be found under the directory *scripts_n_notebooks/vpod_ML_workflows/mine_n_match/data_sources/lmax/VPOD_in_vivo_v1.0_2025-05-09_18-17-49.csv*)

**Genotype Mining:** We queried the NCBI taxonomy database. The standardized NCBI query for opsin coding sequences is detailed in *mine_n_match_functions.ncbi_fetch_opsins*. Mined NCBI data was saved as *mnm_on_all_dbs_ncbi_q_data_cleaned.csv* and *mined_mnm_on_all_dbs_cleaned.fasta.* Instances where we received opsin data from species not part of the query were marked as unintended hits and isolated them for future manual inspection. This data was saved to *mnm_on_all_dbs_ncbi_q_potential_hits.csv*, and species which yielded no opsin data from our queries were saved to *species_w_no_hits.txt*. Merged NCBI and accessory data were saved to *VPOD_in_vivo_v1.0.csv* and *VPOD_in_vivo_v1.0.fasta*. All data from this process are under the report directory *scripts_n_notebooks/vpod_ML_workflows/mine_n_match/mnm_data/mnm_on_all_dbs_2025-02-24_16-29-54* in VPOD.

**Matching Process:** Sequences flagged by OPTICS as '*blastp unsuccessful'* were dropped before matching. Sequences with 100% identity to existing VPOD entries or with a difference >15nm between predicted and *in-vivo* $\lambda_{max}$ were dropped after matching. The final MNM dataframe was saved as *mnm_on_vpod_in_vivo_results_fully_filtered.* The MNM workflow, *mine_n_match-wf.ipynb*, is under the directory *scripts_n_notebooks/vpod_ML_workflows/mine_n_match* in VPOD.

#### S7. *VPOD_v1.3* Integration and *OPTICS_v1.3* Release

*VPOD_v1.3* sequences and metadata were formatted as per Frazer et al. (2024). The five new models (*whole-dataset-mnm*, *wildtype-mnm*, *vertebrate-mnm*, *wildtype-vert-mnm*, *invertebrate-mnm*) and their bootstrap versions are available in *OPTICS_v1.3* on the OPTICS GitHub repository (https://github.com/VisualPhysiologyDB/optics).
